# Unmixing of Imaging Mass Spectrometry Measurements Using Microscopy-Informed Constraints

**DOI:** 10.1101/2025.10.29.685281

**Authors:** R.A.R. Moens, N.H. Patterson, L.G. Migas, A.B. Esselman-Lawrence, F.A. Moser, J.M. Spraggins, R. Van de Plas

## Abstract

Imaging mass spectrometry (IMS) provides spatially resolved molecular information of organic tissue but can be limited by pixel signals mixing contributions from adjacent biological structures, *e.g*. of single cells and multicellular functional tissue units (FTUs). This paper proposes computational methods to predict mass spectral profiles of biological structures on the basis of IMS data by “unmixing” pixel-level signals, leveraging microscopy-based boundary information of these structures. By modeling each biological structure as having a unique mass spectrum, we formulate a linear mixing model and solve the corresponding inverse problem that unmixes blended signals. In particular, we cover both overdetermined and underdetermined linear system scenarios and compare ordinary least squares, nonnegative least squares, and singular value thresholding to a custom algorithm, coined Tissue-informed Unmixing of Labeled regions by Inverse Problem (TULIP), specifically tailored to IMS data. Validation on a synthetic *in-situ* single cell dataset and demonstration on a largescale kidney FTU dataset illustrate the potential of these methods for enhanced *in-situ* tissue structure analysis, e.g. in cellular and tissue studies.

## I. Introduction

Imaging mass spectrometry (IMS) has emerged as a powerful tool for spatially resolved, label-free molecular analysis across a range of biological samples, from large tissue sections to individual cells [1]–[6]. It enables *in-situ* molecular mapping of proteins, lipids, and metabolites, without the need for chemical stains or a priori information [1], [2], making it valuable for investigating biological processes such as disease pathology, tissue heterogeneity, and metabolic changes [7]–[9]. Although there are various surface sampling approaches, MALDI is one of the most common due to its combination of high spatial fidelity and molecular coverage.

A commonly encountered issue within MALDI, with potentially significant impact, is the imperfect spatial alignment between the sampling pattern, here simplified as a measured IMS pixel, and the underlying biological structures. Many structures or regions of interest, such as organelles (depending on the type, below ≈ 1 µm), small-scale cells (depending on the type, at around or below 5 *µ*m), groups of cells (depending on the type and number, at or above 5 *µ*m), or larger-scale functional tissue units (FTUs, depending on the type ≈ 100 µm), may be smaller than the pixel size (at or around 5 µm) and/or span across neighboring pixels, leading to blended or mixed spectra [10]– [12]. This blending limits the specificity of molecular assignments to these structures or regions of interests, particularly in spatially heterogeneous contexts. While instrumental advances like oversampling [13] and transmission geometry MALDI [14], [15] have pushed pixel sizes downwards as a partial solution, they often compromise throughput, sensitivity, or molecular coverage.

Several computational approaches have sought to resolve these limitations by unmixing coarse IMS pixels into regions of interest using prior information, such as provided by microscopy modalities [10], [16]–[19]. They offer higher spatial resolution but most of the time present either insufficiently powerful (local) weighing schemes, are tightly coupled to a specific instrumental resolution, rely on physical cell isolation, or are not easily generalized across tissues or imaging modalities. To address these, we propose TULIP (Tissue-informed Unmixing of Labeled regions by Inverse Problem), a framework that aims to predict the molecular profiles of structures of interest from IMS data by using microscopy-informed constraints. TULIP formulates spectral unmixing as an inverse problem that incorporates spatial priors, *e*.*g*., from microscopy, without requiring physical isolation of the regions of interest. It combines biologically motivated constraints such as sparsity, non-negativity, and low-rank assumptions [20], [21], and is designed to operate across different spatial scales and biological hierarchies, from (sub)cellular domains to tissue structures to anatomical regions. We take a heuristic approach leveraging nuclear norm minimization as a relaxation of rank minimization [22], and bridge ideas from maximum-margin matrix factorization [23], rewriting the nuclear norm as a non-convex Frobenius norm loss [24] to reduce the memory complexity. Finally, we considered advances in non-negative matrix completion [25], as well as regularized non-negative matrix factorization (NMF) [26] to construct our optimization problem. Although solving our proposed constrained low-rank matrix approximation remains generally NP-hard [27], (non-)convex relaxations and heuristic optimizers have provided a practical path forward for our setting.

## II Methods

### A. Inverse Problem Formulation

Our methodological framework defines the combination of IMS with high spatial resolution microscopy data as a forward model. Each biological structure is modelled as a region of interest (ROI), *e*.*g*., a single cell, a cell type group, or an FTU. Each ROI is assumed to be deterministic and homogeneous in molecular/spectral content, thus represented by a unique mass spectrum. Our model considers IMS pixels to represent spectra as linear mixtures of the underlying ROI profiles, *e*.*g*., weighted proportionally by their spatial overlap with IMS pixels. Formally, we express the forward model as:

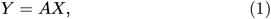

where *Y* ∈ ℝ^*p*×*q*^ represents IMS measurements (known) across p pixels and q mass-to-charge (*m/z*) bins, *A* ∈ ℝ ^*p*×*r*^ is the microscopy-derived spatial mixing matrix (known) indicating overlaps between IMS pixels and annotated ROIs (*e*.*g*., individual cells, cell type groups, or FTUs), and *X* ∈ ℝ^*r*×*q*^ contains the predicted spectra corresponding to each annotated ROI (unknown) with r the total number of such ROIs (known). Note that the mixing matrix A could include a more advanced weighting scheme, incorporating prior knowledge on *e*.*g*., the laser ablation spot morphology or accounting for a non-uniform grid of pixels. Finally, the heterogeneity of the samples, the quality of annotations in A, and available computational resources could also be considered when constructing the model and selecting the right solution method.

#### 1) Single Cell Example

To account for non-annotated regions (NARs, see **Fig. 1**), arising *e*.*g*., due to the morphology of the tissue, we extend the formulation in Eq. 1 by:

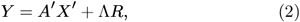

where *Y* ∈ ℝ^*p*×*q*^ represents IMS measurements, *A*′ ∈ ℝ ^*p*×*r*^ is the microscopy-derived spatial mixing matrix, *X*′ ∈ ℝ^*r*×*q*^ contains the predicted ROIs (here single cells spectra), Λ ∈ ℝ ^*p*×*k*^ captures residual spatial weights for k non-annotated regions ensuring that rows of A (see Eq. 3) sum to unity^1^, and *R* ∈ ℝ ^*k*×*q*^ represents the estimated spectra for these nonannotated regions. Note that we model the NAR in each IMS pixel separately (*k* = *p*), but we could also model it as a single spectrum by setting *k* = 1.

**Fig. 1.**
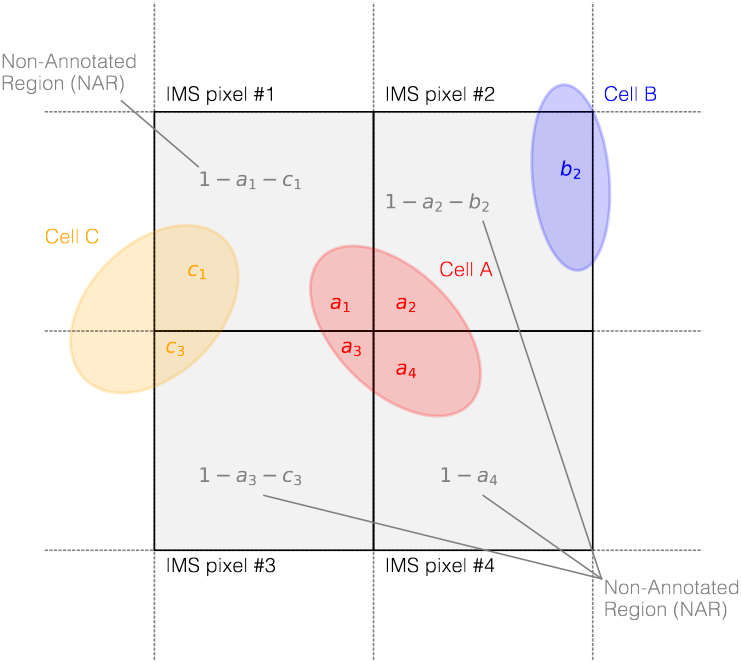
Simplified representation of IMS pixels and annotated ROIs (in this case single cells). Lowercase letters indicate relative overlap areas within an IMS pixel, normalized to unity.

As such,

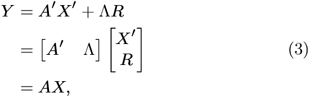

where for the particular example shown in **Fig. 1**

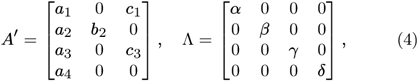

with *α* = 1 − *a*_1_ − *c*_1_, *β* = 1 − *a*_2_ − *b*_2_, *γ* = 1 − *a*_3_ − *c*_3_, and *δ* = 1−*a*_4_. Each *a*_*i*_, *b*_*i*_ and *c*_*i*_ correspond to the relative overlap in area of the ROIs, *i*.*e*. cell, with the particular IMS pixel *i* such that the following holds:

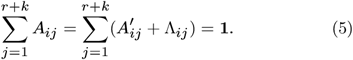

Note that depending on the setting and the modelling, the condition number of the matrix A can be affected by modelling NARs. In turn, this can lead to instabilities if not properly addressed by the method.

### B. Linear System

Given the data sizes typical of these IMS problems, where matrices with more than 100, 000 rows and columns are not uncommon [20], we opt for a straightforward and efficient linear system approach. Commonly, two scenarios are encountered in such an approach:

#### 1) Overdetermined Linear System

This scenario encompasses situations where the mixing matrix A is tall and narrow, *i*.*e*., more rows than columns. For example, it can, but not exclusively, consists of setups in which non-annotated regions (NARs) are either absent or only partially present, and/or where ROIs are relatively large with respect to the IMS pixel size. Following the notation from Eq. 1, this case is described by:

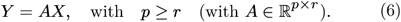

Usually, no exact solution exists^2^. However, a unique least squares solutions can exist^3^ and be efficiently obtained. While computationally efficient, the least squares approach may *e*.*g*., not fully capture detailed residual contributions in highly spatially heterogeneous samples, such as the NAR in the single cell example of **Fig. 1**.

#### 2) Underdetermined Linear System

This scenario applies where the mixing matrix A is short and wide, *i*.*e*., more columns than rows. For example, it can, but not exclusively consists of setups where the comprehensive contributions from NARs or multiple annotated regions per single IMS pixels are modelled. In this case, again following the notation from Eq. 1, the number of unknowns exceeds the number of equations, this case is described by:

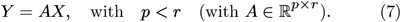

Because the system has more unknowns than equations, additional constraints or prior knowledge, *e*.*g*., regularization or sparsity constraints, are necessary to ensure uniqueness of a solution. Although often more computationally demanding to solve, this model allows for a detailed representation, enabling more rigorous downstream analyses and possibly increasing model fit of the NAR and ROI spectra. The overdetermined model is preferable when residual contributions are thought to be minimal or computational efficiency is prioritized, whereas the underdetermined model can be useful, *e*.*g*., for capturing substantial non-annotated variability.

### C. Prior Knowledge on IMS and Microscopy Data

In many problems involving IMS data, leveraging relevant prior knowledge can significantly improve the accuracy, robustness, and interpretability of the results by constraining the solution space [20], [21], but it can also improve the computational and memory complexity. Examples of such prior information include:

#### 1) Measurement matrix *Y*

- Measured IMS data in *Y* are sometimes subjected to a non-linear transformation f(·), *e*.*g*., due to detector clipping, thresholding measured ion intensity values *M*_*ij*_ that fall under a particular threshold *ϵ* ∈ ℝ_+_ [20]:

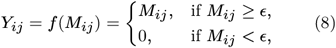

resulting in sparsity levels of the full profile IMS data below 10%.

- Contains a measurement error, *e*.*g*., originating from sample preparation procedures and instrumental artifacts (*e*.*g*., delocalization, matrix-solution binding, laser pattern deformation, detector jitter) assumed to be distributed in a Gaussian fashion.
- High-dimensional: commonly more than 200, 000 *m/z*-bins per pixel, and usually more than 100, 000 pixels.

#### 2) Mixing matrix *A*

- Usually very high sparsity,
- Full-rank, possibly high condition number,
- Bounded, *i*.*e*., ∥*A*∥_∞_ ≤ 1 with entries 0 ≤ *A*_*ij*_ ≤ 1,
- Possibly disturbed by an error term Δ*A, i*.*e. A*+Δ*A*.

#### 3) ROI spectra *X*

- Consists of sums of dependent and independent Poisson-distributed variables,
- Spectra measured independently, but *m/z*-bins exhibit interdependencies,
- Exhibits low-rank and sparsity characteristics.

Using this information and knowing that measured ion intensities are non-negative values, we propose a novel method for solving inverse problems occurring in IMS by using microscopy-informed constraints.

### D. Tissue-informed Unmixing of Labeled regions by Inverse Problem (TULIP)

We propose the TULIP method, incorporating prior knowledge and related to regularized least squares with rank, sparsity and non-negativity constraints.

#### 1) Proposed Program

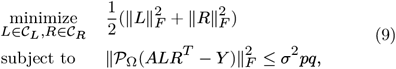

where

- 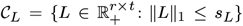 with *t* denoting the factorization rank, and non-negativity promoting biological interpretability;
- 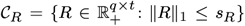 with *t* denoting the factorization rank, and non-negativity promoting biological interpretability;
- s_*L*_, s_*R*_ are sparsity parameters that limit the *ℓ*_1_-norm of *L* and *R* respectively, promoting solution uniqueness and alignment to physical constraints and thus interpretability [21], [28];
- 𝒫_Ω_ is the projection operator onto the set of observed indices Ω, *i*.*e*., for any matrix *Z* ∈ ℝ^*m*×*n*^,
- *σ* represents the standard deviation of the measurement noise (under the assumption that it is modelled by a Gaussian distribution), so that the constraint 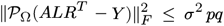 limits the reconstruction error to a level consistent with the noise floor;
- the term *pq* corresponds to the total number of elements in *Y* ∈ ℝ^*p*×*q*^, ensuring that the error tolerance scales with the size of the problem;
- 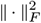 is the squared Frobenius norm.

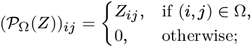

TULIP builds on a low-rank approach based on a nuclear norm minimization [22], a proxy for the non-convex rank minimization, and makes use of concepts from maximum-margin matrix factorization [23] to obtain the Burer-Monteiro approximation [24]. For the latter, it has been proven, under specific conditions, that such non-convex formulations can avoid spurious local minima and recover global optima in polynomial time, see *e*.*g*., [29]. However, we will not assume any of these conditions to hold for our proposed problem. Merely, its explicit factorization of *X* = *LR*^*T*^ simplifies our large-scale optimization and reduces the memory complexity. In addition, parallels between our proposed program and a regularized nonnegative matrix factorization [26], [30]–[32] and non-negative matrix completion can be drawn [21], [25]. Our approach is purely heuristic, combining elements of (non-negative) matrix completion and low-rank matrix approximation together with prior information to provide a practical pathway for addressing our problem in its high-dimensional biological setting. Note that we consider the completion projector 𝒫_Ω_ as a practical construct, as the sampling pattern in IMS is often intensity-based. This means that low intensity values are removed, leading to non-uniform sampling schemes and thus typically no adherence to incoherence or restricted isometry properties [20]. The formulation including the projector 𝒫_Ω_ allows, however, for the sparse format of *Y* and *ALR*^*T*^ to be efficiently exploited and reduce the memory footprint, as no dense matrix of size *p* × *q* is required in memory during optimization.

#### 2) Projected Gradient Descent

Given the inherent sparsity and scale of the data, we employ a projected gradient descent with Adam optimizer (for a parameter discussion see Appendix IV-D) to set adaptive learning rates [33]. The primary advantages of this approach are its flexibility, simplicity, and efficient memory usage without the need for matrix inverses. An outline of the optimization is given in **Algorithm 1**, where the projection onto *𝒞*_*e*_ = {*Q* : *Q* ≥ 0, ∥*Q*∥_1_ ≤ *s*_*e*_} is performed by a soft-thresholding with threshold *λ*_*e*_. We choose to check the fidelity constraint 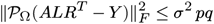 at the final iteration, but this can also be done during the optimization.

##### Algorithm 1

Projected Gradient Descent for TULIP

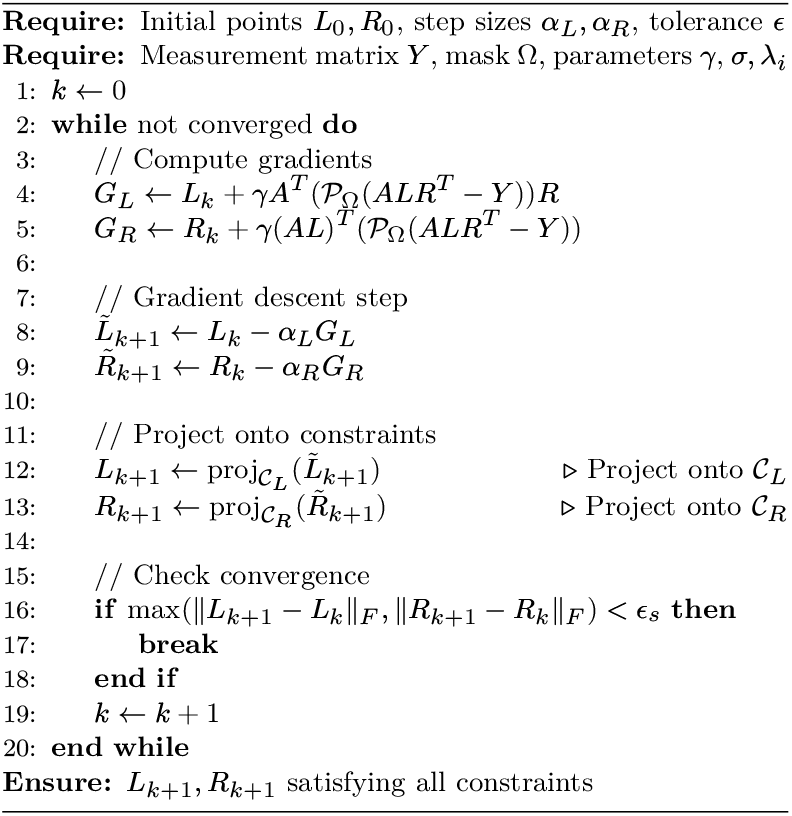

To prevent dense reconstruction of the matrix 𝒫_Ω_(*ALR*^*T*^ *− Y*) when computing the gradients, we implemented a sparse error calculation method leveraging Numba’s just-in-time (JIT) compilation and CPU parallelization. Specifically, we compute *ALR*^*T*^ *− Y* only at the known (observed) indices, Ω, provided by the sparse representation of the matrix *Y*, to significantly reduce memory consumption and computational complexity. Future strategies to expand to even larger datasets could make use of a distributed framework (see *e*.*g*., [34]). However, for our problem we opted for an in-memory approach.

##### a) Computational Complexity Analysis

The primary computational cost per iteration in Algorithm 1 stems from matrix multiplications and sparse matrix operations involved in the gradient computations. Specifically, the approximation of the product 𝒫_Ω_(*ALR*^*T*^ *− Y* ) occurs in two stages. Initially, the intermediate product *AL* is calculated with a complexity of nnz(*A*) · *t*, where *A* ∈ ℝ^*p*×*r*^ is a sparse matrix and nnz(*A*) denotes the number of computed non-zero elements stored in *A*. Subsequently, each column of *R*^*T*^ is individually multiplied by selected rows from the intermediate product, according to a given sparsity pattern defined by the sparse data structure. This step has a complexity proportional to the number of nonzero elements, specifically nnz(*Y* ) · *t*. This step is parallelized, but remains a matrix-vector operation (BLAS level 2) in our implementation. Finally, the computed intermediate results are subtracted from the known *Y*, which is negligible in computational cost. The gradient calculation for *L* involves computing:

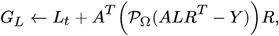

which has two major computational costs. First, computing the residual 𝒫_Ω_(*ALR*^*T*^ −*Y* ) costs (nnz(*A*) +nnz(*Y*)) · *t* operations. Then, multiplying this sparse residual by *R* incurs an additional cost of nnz(*Y* ) · *t*, and subsequently multiplying by A^*T*^ is in the worst case bounded by *rpt*. An analogous complexity can be derived to compute the gradient *G*_*R*_. An overview with comparison to an all dense complexity is given in **Table I**. Generally speaking, TULIP is of computation complexity 𝒪(*prt*). This comparison underscores the efficiency of the algorithm when handling large, sparse matrices and highlights its scalability given that the latent dimensions r and t are mostly of modest size relative to the overall problem dimensions *p* and *q*. Of course, the algorithm is practically limited by the number of BLAS level 2 operations in the calculation of the error term 𝒫_Ω_(*ALR*^*T*^ *− Y*), which will be our bottleneck. Finally, note that this is the per iteration complexity.

**TABLE I.**
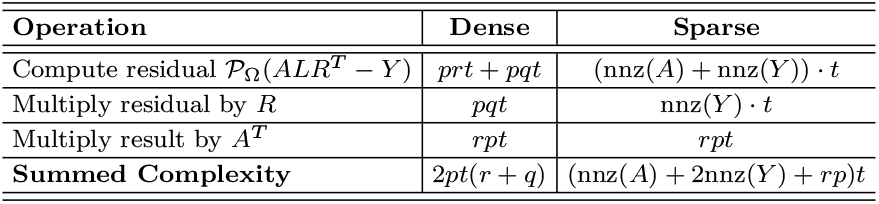
Per iteration computational complexity comparison between dense and sparse matrix format of ***Y*** AND ***A*** for computing the gradient of ***L, G***_***L***_ **→ *L***_***t***_ **+ *A***^***T***^ (***𝒫***_Ω_**(*ALR***^***T***^ **− *Y* ) )*R***.

##### b) Memory Complexity Analysis

The memory cost for *Y* is on the order of 𝒪(nnz(*Y*)). The same nnz(*Y*) is true for P_Ω_(*ALR*^*T*^ −Y). The matrix A ∈ ℝ^*p*×*r*^ is a sparse matrix with associated cost 𝒪(nnz(A)). The low-rank factors *L* and *R* are defined by the factorization rank t. The memory needed to store these matrices is 𝒪(*rt*) for *L* and 𝒪(*qt*) for *R*. The same costs are required for the gradients and an additional copy needs to be stored from the previous time step *k*. Collecting all the storage costs, we have a memory complexity:

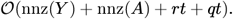

Since we assume that the size t is much smaller than *p, q*, and *r* (*i*.*e*., t ≪ min(*p, q, r*)) and that the number of nonzeros in *Y* satisfies nnz(*Y* ) ≪ *pq*, and the nnz(*A*) ≪ nnz(*Y* ), the following simplifications hold:

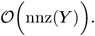

This expression indicates that, in our sparse and low-rank setting, the memory required is much lower than that of a dense formulation, which would be of the order 𝒪(pq). This also emphasizes the importance of the efficient exploitation of the projection operator 𝒫_Ω_. Practically, we observe a peak memory footprint of ∼ 3 times the data footprint of *Y*, assumed to be represented as a sparse matrix.

##### c) Parameter Optimization

We have the following parameters in our optimization:

- *γ* ∈ ℝ_+_: defines the trade-off between smoothness of *L* and *R*/low-rank property^4^ and the data fidelity constraint;
- *t* ∈ ℕ: factorization rank, with the intuition to set it higher than the true underlying rank [29], also plays an important role in the convergence rate and computational cost per step (see **Table I**);
- *σ* ∈ ℝ_+_: standard deviation of the noise in the measurements, can be estimated *a priori*;
- *ϵ*_*s*_ ∈ ℝ_+_: stopping criterion;
- *λ*_*L*_ ∈ ℝ_+_: element-wise soft-threshold for L, influences the sparsity pattern of L;
- *λ*_*R*_ ∈ ℝ_+_: element-wise soft-threshold for R, influences the sparsity pattern of R.

**TABLE II.** Comparative Performance Metrics of LS, NNLS, SVT and TULIP methods in an overdetermined linear system setting for a raw, full profile (**415 007** features) dataset. Metrics include reconstruction error, cell fit score, non-negativity, sparsity, and clustering score, with computational performance assessed through run times (see Appendix IV-E). Methodology validated via **100**-fold bootstrapping using **10 000** ims pixels and **500** *m/z*-bins with Gaussian-weighted mixing (see Appendix IV-A).

| Performance Metrics - Overdetermined Scenario |  |  |  |  |  |  |
| --- | --- | --- | --- | --- | --- | --- |
| Method | Recon. Error (%) | Cell Fit Score (%) | Non-Negativity (%) | Sparsity (%) | Clustering Score (%) | Run Time (s) |
| LS | 24.8540<sup>+4.8861</sup><sub>-8.2822</sub> | 75.0359<sup>+11.5675</sup><sub>-22.6169</sub> | 90.7511<sup>+0.9268</sup><sub>-0.9540</sub> | 9.3480<sup>+0.8589</sup><sub>-0.8602</sub> | 85.2260<sup>+10.3246</sup><sub>-28.2804</sub> | 0.0434<sup>+0.3193</sup><sub>-0.0226</sub> |
| NNLS | 24.8691<sup>+4.8834</sup><sub>-8.2837</sub> | 80.5795<sup>+7.6214</sup><sub>-24.1341</sub> | 100.0000<sup>+0.0000</sup><sub>-0.0000</sub> | 10.0708<sup>+0.8425</sup><sub>-1.0307</sub> | 91.8206<sup>+4.0296</sup><sub>-24.4886</sub> | 3.2044<sup>+0.4922</sup><sub>-0.4969</sub> |
| SVT | 24.8562<sup>+4.8910</sup><sub>-8.2819</sub> | 70.5367<sup>+15.4892</sup><sub>-53.4948</sub> | 90.5926<sup>+0.8669</sup><sub>-0.9769</sub> | 9.4617<sup>+0.9449</sup><sub>-0.8477</sub> | 79.1175<sup>+16.3560</sup><sub>-27.0554</sub> | 7.7211<sup>+2.0974</sup><sub>-1.8037</sub> |
| TULIP | 25.3831<sup>+4.8147</sup><sub>-7.7523</sub> | 80.9436<sup>+6.3865</sup><sub>-28.2552</sub> | 100.0000<sup>+0.0000</sup><sub>-0.0000</sub> | 9.1826<sup>+2.6137</sup><sub>-2.4993</sub> | 87.9596<sup>+7.3921</sup><sub>-29.6487</sub> | 6.9157<sup>+0.3808</sup><sub>-0.3025</sub> |

In practice, the most important parameters to adjust are *γ, t, λ*_*L*_, and *λ*_*R*_. For *λ*_*L*_ and *λ*_*R*_ the intuition is straightforward: increasing these parameters leads to increased sparsity in, respectively, *L* and *R*. On the other hand, *γ* and t will play a role in the reconstruction error (see Appendix IV-E). The Ordinary Least Squares (see Section II-D.3) solution can usually provide

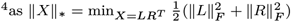

a lower bound for this reconstruction error. We apply data subsampling and a Bayesian parameter optimization over *γ, t, λ*_*L*_, and *λ*_*R*_ to speed up the parameter tuning. The parameter tuning per case study is presented in Appendix IV-D.

#### 3) Benchmarking Methods

For benchmarking our novel method, we used three “off-the-shelf” methods: Ordinary Least Squares (LS), Non-negative Least Squares (NNLS), and Singular Value Thresholding (SVT) [35]. The ordinary least squares is implemented through an LU decomposition exploiting the sparse matrices [36], while the NNLS is implemented through an alternating direction method of multipliers (ADMM) [37]. Finally, the singular value thresholding is making use of the GESDD singular value decomposition implementation.

- **Ordinary Least Squares (LS):**

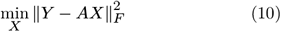

- **Non-Negative Least Squares (NNLS):**

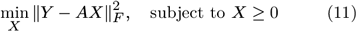

- **Singular Value Thresholding (SVT):**

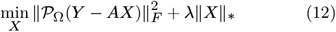

A nice property of LS is that it will theoretically provide us a lower bound for both the NNLS, SVT, as well as TULIP, when considering the metric 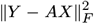. Note that LS and NNLS do not include completion of missing values, but can still be useful when assuming *Y*_*ij*_ = *M*_*ij*_, *i*.*e*., missing values were assumed to be zero. Reciprocally, for SVT and TULIP, we can assume all values to be measured. As such, any of these methods can be run on either sparse raw (non-feature-selected) data or on dense peak-picked (feature-selected) data. In general, the overall computational complexity of LS will be considered here as 𝒪(*pr*^2^) [38], similar to NNLS as 𝒪(*pr*^2^), and the complexity of the SVT as 𝒪(*pqt*). All methods, including TULIP, can be used both in overdetermined as well as underdetermined scenarios. However, note that the properties of the solution will differ.

#### 4) Availability

The implementations of the methods used in this paper, are provided as an open-source Python 3 toolbox at https://github.com/vandeplaslab/tulip.

## III. Numerical Results

The first numerical study provides a ground truth comparison for both under- and overdetermined scenarios, the second study allows us to investigate the algorithm’s performance in a large-scale setting. For the first study, synthetic single cell datasets were generated by spatial segmentation of multiplexed immunofluorescence (MxIF) microscopy images. IMS spectra for the single cells were simulated using *k*-means clustering of the corresponding IMS spectra (obtained after the MxIF imaging) to obtain ground truth spectra. The downsampling of microscopy data to IMS resolutions (0.65 µm to 5 µm)was performed linearly with a Gaussian-weighting. Methods were validated through 100-fold bootstrapping, assessing reconstruction accuracy (*i*.*e*., optimization fit), cell fit scores (*i*.*e*., ground truth fit), sparsity (*i*.*e*., indicator for sparsity in the solution space), clustering quality (*i*.*e*., closeness to ground truth classification), and computational efficiency. For further details on the synthetic data, statistical resampling and metrics, we refer the reader to the Appendix. A large-scale FTU dataset (details in Appendix), previously employed in a study [39] and part of the Human BioMolecular Atlas Program (HuBMAP) study of Human Kidney [40], was integrated into our analysis for the second study. The parameter setting for both studies can likewise be found in the Appendix. All calculations were performed on CPU on a Dell Precision 7920 workstation equipped with 56 cores at 2.7 GHz and 1.5 TB of memory.

### A. Synthetic Single Cell

By generating ground-truth spectra through clustering real IMS spectra and mixing those, we do not ensure that they replicate the sparsity patterns inherent to the raw IMS data matrix *Y*. This is no problem as we can define Ω accordingly and prove the applicability of our methodology on both clipped and non-clipped data. At the same time, note that we do not optimize for *X* to be sparse, but rather *L* and/or *R* for optimization reasons.

#### 1) Overdetermined Linear System

In the overdetermined case we modelled the NAR using a single spectrum. We subsampled the raw IMS spectra to create our ground truth spectra and varied the underlying spatial cell distribution by altering the section used of the microscopy image to different spatial locations to construct the mixing matrix A. The results are presented in Table II. We observed that all methods achieved comparable reconstruction errors (*i*.*e*., optimization error). TULIP exhibited a slightly higher reconstruction error, which is expected due to the nature of its gradient descent optimization strategy. The cell fit scores (*i*.*e*., ground truth fit) revealed that the nonnegative constrained methods (NNLS and TULIP) performed approximately 5% better than LS and 10% better than SVT. This suggests that enforcing non-negativity enhances biological interpretability and alignment with known ground truth distributions. Notably, TULIP achieved the highest average cell fit score, implying better recovery of biologically relevant spatial structures despite its slightly elevated reconstruction error. We also observed a substantially larger confidence interval for SVT in both the cell fit and clustering scores. This variability is caused by the problem becoming ill-conditioned for some spatial settings. And although that we regularize both the LS and SVT by a small factor, it seems to affect the SVT more than the LS. Sparsity was observed similar across all methods, as expected, since it was not considered a central concern in this case study. This confirms that none of the methods excessively pruned the solution space under this setup. Clustering scores, while generally decent, showed greater variability, especially for SVT. This metric is merely used here, however, to demonstrate a post-processing capability, rather than a dedicated clustering. Nonetheless, consistent clustering scores across methods support the robustness of spectral decomposition approaches in preserving cell-type distinctions. As such, it messages that a good clustering can be obtained from the estimated spectra provided by the different methods. Finally, run times varied based on both the number of iterations required by each method and their implementation specifics. While this metric only provides a rough estimate, it is evident that LS is the most computationally efficient, followed by NNLS, TULIP, and SVT, which incurs the highest computational cost.

By application of a similar bootstrap method (see Appendix) on the LS data, uncertainty intervals were obtained for the reconstruction of individual cell spectra (see **Fig. 2**). We observe there that the confidence interval is small with respect to the relative intensities for this particular class. In practice, this might change from cell to cell and depends heavily on the setting, *e*.*g*. conditioning of the A matrix. Note, that this uncertainty is related to the mixing process, other sources of uncertainties are not considered, but could impact the overall uncertainty.

**Fig. 2.**
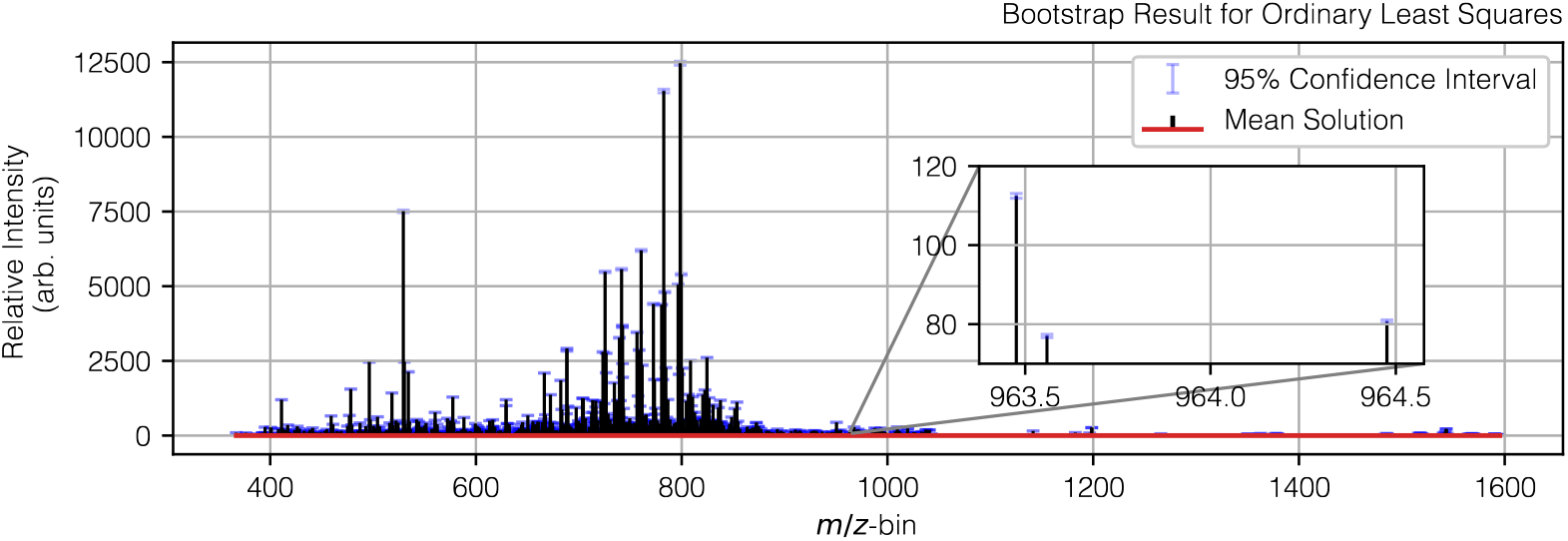
Bootstrap results for ordinary least squares estimates in the overdetermined scenario, showing the result for an individual retrieved ROI, *i*.*e*., cell, spectrum and its corresponding confidence interval. One observes that the confidence interval is relatively small with respect to the relative intensities for this particular single cell, even in the inset example on the right. This will mainly depend on the conditioning of the ***A*** matrix. Similar plots are obtainable for the other methods.

#### 2) Underdetermined Linear System

In the underdetermined case (see **Table III**), where each NAR is modelled as an individual spectrum, all methods achieved a low reconstruction error overall as well. As there are more degrees-of-freedom to the model, the reconstruction error is remarkably smaller than the overdetermined linear system. TULIP has slighly higher reconstruction error, as previously observed in the overdetermined case. This is thought to originate from slow convergence in the neighborhood of its local minimum. In terms of cell fit score NNLS performs best, suggesting improved biological relevance. As in the overdetermined case, non-negativity appears crucial: TULIP and NNLS strictly enforces this constraint, unlike LS and SVT, and this likely contributes to a superior interpretability. The sparsity is, however, notably higher in LS and SVT, and consistently lower for TULIP. However, since the latter is mainly driven by parameter tuning, it has the potential to be improved at the cost of a higher reconstruction error. Clustering scores were comparable, with NNLS slightly ahead, yet TULIP remained robust across trials. While TULIP incurred substantially longer run times—2 to 6 times slower than SVT, this trade-off is balanced by its gradient-based nature and better scalability in high-dimensional settings without relying on the SVD. By application of a similar bootstrap method for TULIP, uncertainty intervals are obtained for the individual cell spectra (see **Fig. 3**).

**Fig. 3.**
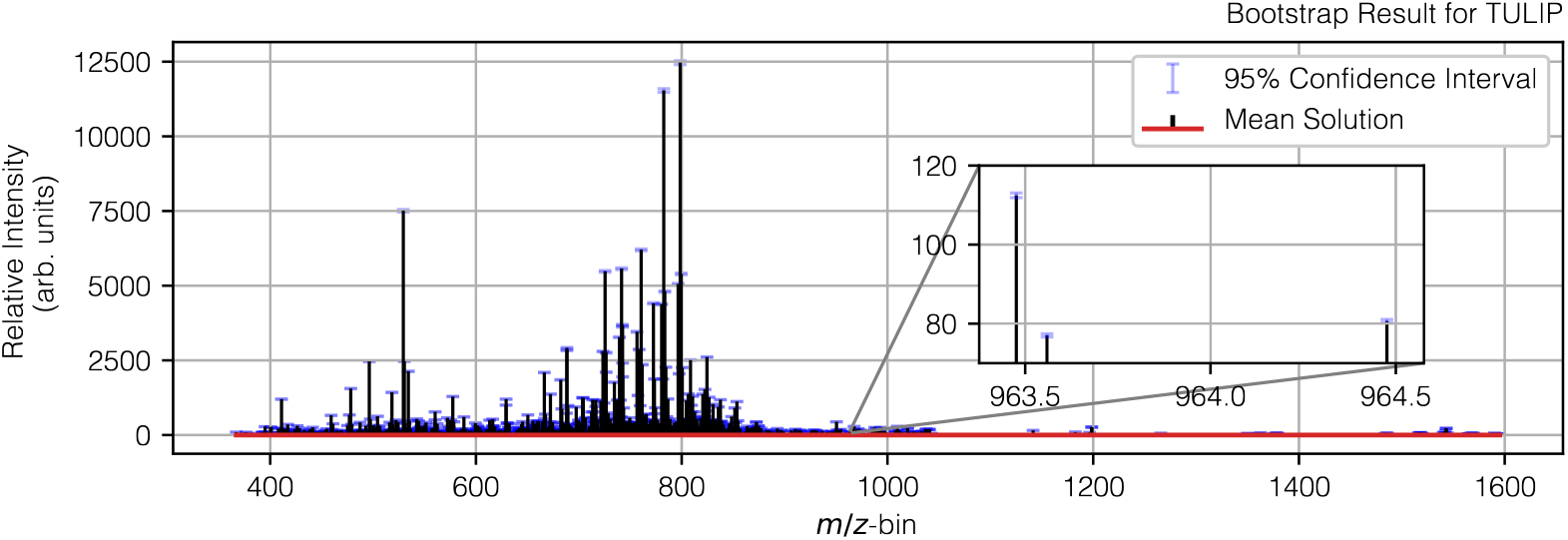
Bootstrap results for TULIP in the underdetermined scenario, showing the result for an individual retrieved cell spectrum and its corresponding confidence interval. One observes that the confidence interval is relatively small with respect to the relative intensities for this particular single cell, even in the inset example on the right, and comparable to LS, see Fig. 2. This will mainly depend on the conditioning of the ***A*** matrix. Similar plots are obtainable for the other methods.

**TABLE III.** Comparative Performance Metrics of ls, NNLS, SVT and TULIP methods in an underdetermined linear system setting for a raw, full profile (**415 007** features) dataset. Metrics include reconstruction error, cell fit score, non-negativity, sparsity, and clustering score, with computational performance assessed through run times (see Appendix IV-E). Methodology validated via **100**-fold bootstrapping using **10 000** ims pixels and **500** *m/z*-bins with Gaussian-weighted mixing (see Appendix IV-A).

| Performance Metrics - Underdetermined Scenario |  |  |  |  |  |  |
| --- | --- | --- | --- | --- | --- | --- |
| Method | Recon. Error (%) | Cell Fit Score (%) | Non-Negativity (%) | Sparsity (%) | Clustering Score (%) | Run Time (s) |
| LS | 0.2771<sup>+0.0916</sup><sub>-0.0545</sub> | 86.6961<sup>+4.6787</sup><sub>-9.6768</sub> | 83.8352<sup>+2.2559</sup><sub>-2.4131</sub> | 18.5638<sup>+2.5923</sup><sub>-2.5577</sub> | 92.8343<sup>+3.0519</sup><sub>-9.6459</sub> | 0.0705<sup>+0.2082</sup><sub>-0.0233</sub> |
| NNLS | 0.2406<sup>+0.1021</sup><sub>-0.0898</sub> | 88.4062<sup>+4.3031</sup><sub>-10.7151</sub> | 100.0000<sup>+0.0000</sup><sub>-0.0000</sub> | 12.1890<sup>+1.5724</sup><sub>-1.5943</sub> | 94.0690<sup>+2.5314</sup><sub>-4.4621</sub> | 12.9890<sup>+4.1889</sup><sub>-2.3521</sub> |
| SVT | 0.0057<sup>+0.0036</sup><sub>-0.0024</sub> | 86.8179<sup>+4.6750</sup><sub>-9.7444</sub> | 82.8575<sup>+2.2767</sup><sub>-2.4731</sub> | 18.4844<sup>+2.9039</sup><sub>-2.7507</sub> | 93.1346<sup>+2.9369</sup><sub>-9.8467</sub> | 3.6418<sup>+0.9488</sup><sub>-0.4707</sub> |
| TULIP | 2.0967<sup>+0.2924</sup><sub>-0.2380</sub> | 85.2511<sup>+4.6268</sup><sub>-9.8960</sub> | 100.0000<sup>+0.0000</sup><sub>-0.0000</sub> | 7.4594<sup>+2.5942</sup><sub>-2.1238</sub> | 89.5560<sup>+5.5831</sup><sub>-4.5504</sub> | 86.7397<sup>+27.8654</sup><sub>-6.2593</sub> |

### B. HuBMAP Study on Human Kidney (FTU)

To demonstrate the scalability of our algorithm and validate it using non-synthetic data, we conducted a numerical experiment on an IMS dataset [39], *Y* ∈ ℝ^869,851×507,429^, stored in Compressed Sparse Column (CSC) format with a memory footprint of 124.24 GB (equivalent to 1765.55 GB in dense format). The associated mixing matrix *A* ∈ ℝ^869,851×(6+503,926)^ is also stored in CSC format, occupying 0.014 GB (or 1753.39 GB in dense format). This setup includes 6 ROIs, corresponding to different clusters (*i*.*e*., different cell types inside the kidney’s glomerulus). A visualization of a single glomerulus is provided in **Fig. 4**, and individual segment/ROI names are listed in **Table IV**, alongside 503, 926 NARs.

**Fig. 4.**
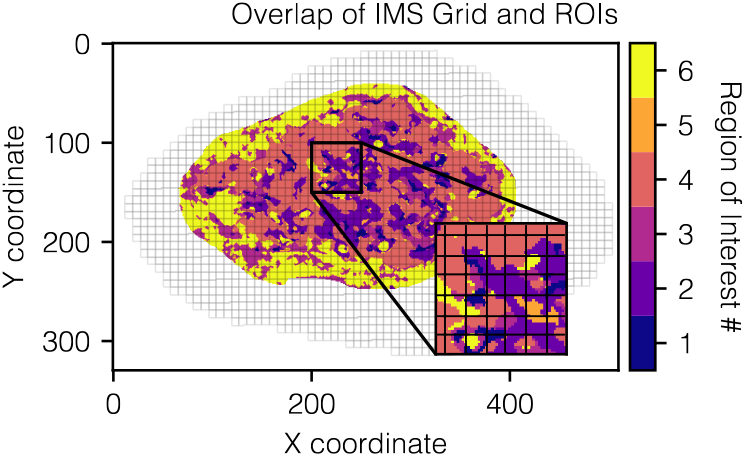
Example of a single glomerulus with highlighted glomerular segments. Observe that the IMS pixel grid overlaps also outside the glomerulus, this padding was added to ensure that the whole glomerulus was covered. The total dataset consists of dozens of these glomeruli. See Table IV for the glomerular segment names.

**TABLE IV.** Glomerular segment naming as seen in Fig. 4. The segments are similar to a previous study [39].

| ROI # | Glomerular Segment |
| --- | --- |
| 1 | Mesangial cells |
| 2 | Endothelial cells |
| 3 | Non-cell specific cluster |
| 4 | Podocytes |
| 5 | Mesangial matrix |
| 6 | Zero/low immunofluorescence staining |

Our implementations of LS and NNLS require an explicit computation of *A*^*T*^*A*, making them infeasible for this dataset. Additionally, the computational complexity of SVT becomes prohibitive and would necessitate either an out-of-memory or a randomized strategy, as previously suggested [20]. SVT also requires computing the pseudo-inverse of the matrix *A*, which is similarly impractical under these conditions and therefore not considered here.

However, TULIP remains applicable: with an average runtime of 37 minutes per iteration over 100 iterations, the total computation time amounted to approximately 60 hours to obtain the following results.

#### 1) Quantitative Analysis

To validate our approach, we compared results with a prior study [39], in which IMS pixels associated with a single cluster (*i*.*e*., no mixing) were averaged per cluster to generate ROI spectra. While one could argue that obtaining spectra in this manner is computationally cheaper, our method offers a scalable alternative suitable for more challenging environments. As such, this analysis validates both the accuracy and scalability of our approach.

**Table V** reports the reconstruction error (see IV-E for more information) across all pixels, as well as for pure (single cluster) and mixed (multiple clusters or cluster/NAR) pixel categories. TULIP significantly reduces the reconstruction error in all categories, with the most substantial improvement observed in the “All” group (from 80.09% to 22.39%), demonstrating advanced overall reconstruction performance. The improvement for “Pure” pixels is smaller, as expected, since these cases are simpler to model (see **Fig. 9** for the reconstruction score per cluster). Additionally, the spectra estimated by TULIP closely match the averaged spectra obtained from pure pixels, lending empirical support to the validity of our method. These average spectra may also serve as effective initializations for TULIP, potentially improving convergence and stability. Finally, substantial gains are observed in the “Mixed” category, further highlighting TULIP’s robustness in handling spectral mixtures. In general, the overall reconstruction error is believed to be constrained by the expressiveness of the mixing matrix *A, i*.*e*. how well A can represent the possible values of Y through linear combinations of *X*. Future work may explore models that explicitly incorporate ΔA to further improve reconstruction accuracy.

**TABLE V.** Reconstruction error comparison between the results used in a previous study [39] and TULIP. For “All”, all pixels (*i*.*e*., **869, 851**) are considered, for “Pure”, only pixels that have a single underlying cluster (*i*.*e*., **66, 367**) are used and for “Mixed”, only pixels having a mix of underlying clusters and clusters and NARs (*i*.*e*. **317, 565**) are used.

| Method | Reconstruction Error (%) |  |  |
| --- | --- | --- | --- |
|  | All | Pure | Mixed |
| Previous Study [39] | 80.09 | 34.89 | 33.18 |
| TULIP | 22.71 | 34.43 | 29.84 |

#### 2) Qualitative Analysis

The spectrum for each individual cluster is shown in **Fig. 7**. While these spectra appear visually similar, a more detailed comparison using the mass spectral difference plots, presented in **Fig. 5**, reveals distinct differences between the spectra of different clusters. This figure is analogous to **Fig. 8**, which was obtained using a procedure similar to that of the previous study [39] (and also mentioned in that study), and thus further supports the validity of our approach.

**Fig. 5.**
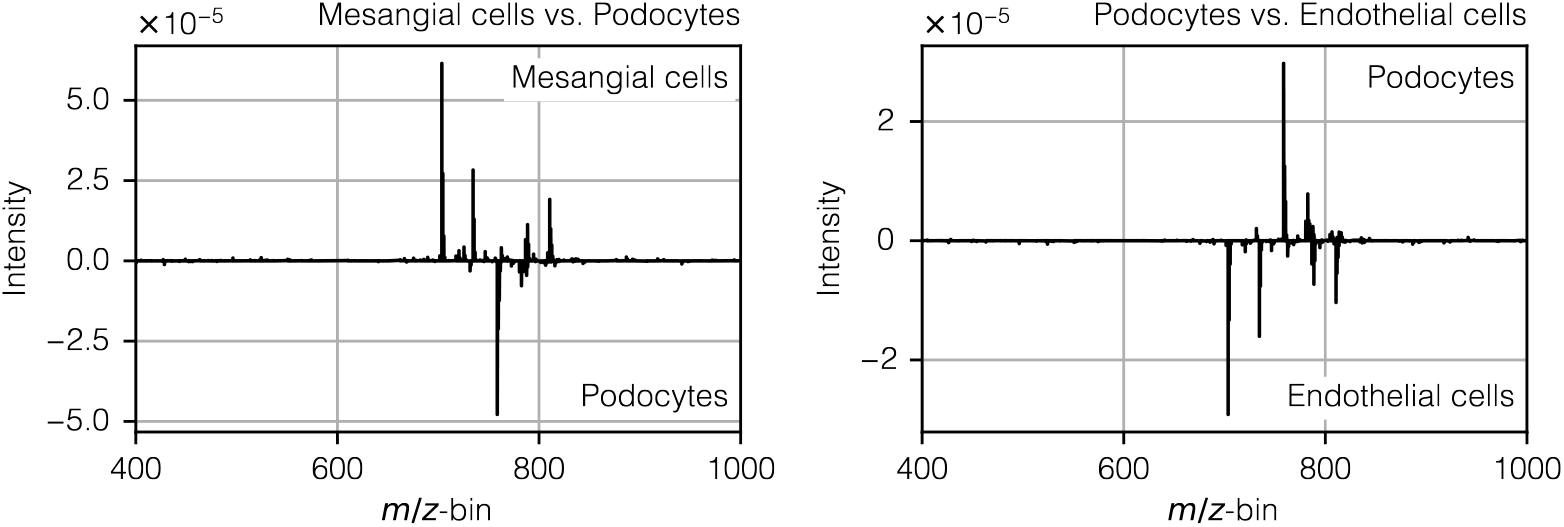
Mass spectral difference (of total sum normalized spectra) plots illustrating ion intensity differences between ROI **#1** (primarily mesangial cells) and ROI **#4** (primarily podocytes) (left), and between ROI **#4** and ROI **#2** (primarily endothelial cells) (right) as obtained by TULIP. These results highlight that each glomerular segment exhibits a distinct profile of IMS-detected molecular species. Comparing these results to Fig. 8 generated with a procedure identical to the previous study [39], we observe that most peaks are further validating our approach.

## IV. Conclusion

This work introduces a novel computational framework that leverages microscopy-informed constraints to overcome a key challenge in IMS: sampling patterns mixing underlying biological structures. We provide a strategy to overcome this limitation in the specificity of molecular assignments at the regions of interest level, particularly interesting in spatially heterogeneous contexts. By formulating spectral unmixing as an inverse problem, our approach effectively disentangles mixed measurements from IMS pixels into the underlying molecular spectra of distinct biological regions, ranging from single cells to larger functional tissue units. Through the integration of biologically meaningful constraints such as non-negativity, sparsity, and low-rank assumptions, a specifically tailored algorithm, TULIP, offers a robust strategy to address both overdetermined and underdetermined scenarios. Comparative studies against established methods, including Ordinary Least Squares (LS), Non-negative Least Squares (NNLS), and Singular Value Thresholding (SVT), demonstrate that, while each method has its merits, TULIP consistently achieves superior cell fit scores and enforces non-negativity in the recovered spectra for the overdetermined case. At the same time, it reports competitive scores and errors to these methods in the underdetermined scenario. These advantages are particularly pronounced in complex settings involving full profile data and significant non-annotated contributions. Furthermore, the evaluation on a challenging large-scale dataset highlights the method’s scalability and its potential to yield biologically interpretable results. Overall, our results underscore the benefit of integrating high-resolution microscopy information to guide the unmixing process in IMS, thereby enhancing molecular specificity and interpretability in tissue imaging. Future work will focus on further refining the parameter tuning process, extending the framework to additional imaging modalities, and addressing challenges posed by expressiveness of the mixing matrix. Ultimately, TULIP paves the way for more detailed *in-situ* analyses of heterogeneous biological samples, opening new avenues for research in cellular and tissue research.

## Acknowledgements

Research reported in this publication was supported by the National Institutes of Health (NIH)’s Common Fund, National Institute Of Diabetes And Digestive And Kidney Diseases (NIDDK), and the Office Of The Director (OD) under Award Numbers U54DK120058, U54DK134302, and U01DK133766 (J.M.S. and R.V.), by NIH’s Common Fund, National Eye Institute, and the Office Of The Director (OD) under Award Number U54EY032442 (J.M.S. and R.V.), by NIH’s National Institute Of Allergy And Infectious Diseases (NIAID) under Award Numbers R01AI138581 and R01AI145992 (J.M.S. and R.V.), by NIH’s National Institute On Aging (NIA) under Award Number R01AG078803 (J.M.S. and R.V.), by NIH’s National Cancer Institute (NCI) under Award Number U01CA294527 (J.M.S. and R.V.), and by the National Science Foundation Major Research Instrument Program CBET – 1828299 (J.M.S.). The research was furthermore made possible in part by grant numbers 2021-240339 and 2022-309518 (L.G.M. and R.V.) from the Chan Zuckerberg Initiative DAF, an advised fund of Silicon Valley Community Foundation. The content is solely the responsibility of the authors and does not necessarily represent the official views of the National Institutes of Health.

## Footnotes

1 In our case, this constraint follows the nature of laser-based ablation, where the same amount of energy is deposited for each IMS pixel surface area.

2 *i.e*., not consistent, *e.g*., when *Y* is not fully in the column space of *A*.

3 Usually when rank(*A*) is full.

## References

[1] R. M. Heeren, D. F. Smith, J. Stauber, B. Kükrer-Kaletas, and L. MacAleese, “Imaging mass spectrometry: Hype or hope?” Journal of the American Society for Mass Spectrometry, vol. 20, no. 6, pp. 1006–1014, 2009.

[2] R. M. Caprioli, “Imaging mass spectrometry: A perspective,” Journal of Biomolecular Techniques: JBT, vol. 30, no. 1, p. 7, 2019.

[3] L. Li, R. W. Garden, and J. V. Sweedler, “Single-cell maldi: A new tool for direct peptide profiling,” Trends in Biotechnology, vol. 18, no. 4, pp. 151–160, 2000.

[4] A. R. Buchberger, K. DeLaney, J. Johnson, and L. Li, “Mass spectrometry imaging: A review of emerging advancements and future insights,” Analytical Chemistry, vol. 90, no. 1, p. 240, 2017.

[5] A. Körber, I. G. Anthony, and R. M. Heeren, “Mass spectrometry imaging,” Analytical Chemistry, 2025.

[6] K. V. Djambazova, J. M. Van Ardenne, and J. M. Spraggins, “Advances in imaging mass spectrometry for biomedical and clinical research,” TrAC Trends in Analytical Chemistry, vol. 169, p. 117344, 2023.

[7] M. Aichler and A. Walch, “Maldi imaging mass spectrometry: Current frontiers and perspectives in pathology research and practice,” Laboratory Investigation, vol. 95, no. 4, pp. 422–431, 2015.

[8] B. Ballú, M. Hanselmann, and R. Heeren, “Mass spectrometry imaging for the investigation of intratumor heterogeneity,” Advances in Cancer Research, vol. 134, pp. 201–230, 2017.

[9] D. Miura, Y. Fujimura, and H. Wariishi, “In situ metabolomic mass spectrometry imaging: Recent advances and difi"culties,” Journal of Proteomics, vol. 75, no. 16, pp. 5052–5060, 2012.

[10] K. Ščupáková, F. Dewez, A. K. Walch, R. M. Heeren, and B. Ballu?, “Morphometric cell classification for single-cell maldi-mass spectrometry imaging,” Angewandte Chemie, vol. 132, no. 40, pp. 17 600–17 603, 2020.

[11] M. Nijs, T. Smets, E. Waelkens, and B. De Moor, “A mathematical comparison of non-negative matrix factorization related methods with practical implications for the analysis of mass spectrometry imaging data,” Rapid Communications in Mass Spectrometry, vol. 35, no. 21, p. e9181, 2021.

[12] P.-L. Delacour, S. Wahls, J. M. Spraggins, L. Migas, and R. Van de Plas, “Signal recovery using a spiked mixture model,” IEEE Transactions on Signal Processing, pp. 1–14, 2025.

[13] A. Maimò-Barceló, J. Garate, J. Bestard-Escalas, R. Fernández, L. Berthold, D. H. Lopez, J. A. Fernández, and G. Barceló-Coblijn, “Confirmation of sub-cellular resolution using oversampling imaging mass spectrometry,” Analytical and Bioanalytical Chemistry, vol. 411, pp. 7935–7941, 2019.

[14] A. Zavalin, E. M. Todd, P. D. Rawhouser, J. Yang, J. L. Norris, and R. M. Caprioli, “Direct imaging of single cells and tissue at sub-cellular spatial resolution using transmission geometry maldi ms,” Journal of Mass Spectrometry, vol. 47, no. 11, pp. 1473–1481, 2012.

[15] R. S. Young, A.-K. Piper, L. McAlary, J. C. McKinnon, J. S. Lum, J. Soltwisch, M. Niehaus, and S. R. Ellis, “Subcellular mass spectrometry imaging of lipids and nucleotides using transmission geometry ambient laser desorption and plasma ionisation,” bioRxiv preprint bioRxiv:2025.05.13.653655, 2025.

[16] T.-H. Ong, D. J. Kissick, E. T. Jansson, T. J. Comi, E. V. Romanova, S. S. Rubakhin, and J. V. Sweedler, “Classification of large cellular populations and discovery of rare cells using single cell matrix-assisted laser desorption/ionization time-of-?ight mass spectrometry,” Analytical Chemistry, vol. 87, no. 14, pp. 7036–7042, 2015.

[17] E. T. Jansson, T. J. Comi, S. S. Rubakhin, and J. V. Sweedler, “Single cell peptide heterogeneity of rat islets of langerhans,” ACS Chemical Biology, vol. 11, no. 9, pp. 2588–2595, 2016.

[18] Y. R. Xie, V. K. Chari, D. C. Castro, R. Grant, S. S. Rubakhin, and J. V. Sweedler, “Data-driven and machine learning-based framework for image-guided single-cell mass spectrometry,” Journal of Proteome Research, vol. 22, no. 2, pp. 491–500, 2023.

[19] L. Rappez, M. Stadler, S. Triana, R. M. Gathungu, K. Ovchinnikova, P. Phapale, M. Heikenwalder, and T. Alexandrov, “Spacem reveals metabolic states of single cells,” Nature Methods, vol. 18, no. 7, pp. 799–805, 2021.

[20] R. A. Moens, L. G. Migas, J. M. Van Ardenne, E. P. Skaar, J. M. Spraggins, and R. Van de Plas, “Preserving full spectrum information in imaging mass spectrometry data reduction,” Bioinformatics, vol. 41, no. 5, p. btaf247, 2025.

[21] F. A. Battjes, K. Olling, R. Van de Plas, and R. A. R. Moens, “Enforcing physical constraints in full spectrum imaging mass spectrometry data reduction,” [Unpublished manuscript], 2025.

[22] M. Fazel, “Matrix rank minimization with applications,” Ph.D. dissertation, PhD thesis, Stanford University, 2002.

[23] N. Srebro, J. Rennie, and T. Jaakkola, “Maximum-margin matrix factorization,” Advances in neural information processing systems, vol. 17, 2004.

[24] S. Burer and R. D. Monteiro, “A nonlinear programming algorithm for solving semidefinite programs via low-rank factorization,” Mathematical Programming, vol. 95, no. 2, pp. 329–357, 2003.

[25] O. V. Thanh and N. Gillis, “Minimum-volume nonnegative matrix completion,” in 2024 32nd European Signal Processing Conference (EUSIPCO). IEEE, 2024, pp. 2452–2456.

[26] L. Taslaman and B. Nilsson, “A framework for regularized nonnegative matrix factorization, with application to the analysis of gene expression data,” PLOS ONE, vol. 7, no. 11, p. e46331, 2012.

[27] N. Gillis, “Introduction to nonnegative matrix factorization,” arXiv Preprint 1703.00663, 2017.

[28] H. Kim and H. Park, “Sparse non-negative matrix factorizations via alternating non-negativity-constrained least squares for microarray data analysis,” Bioinformatics, vol. 23, no. 12, pp. 1495–1502, 2007.

[29] R. Ge, J. D. Lee, and T. Ma, “Matrix completion has no spurious local minimum,” Advances in Neural Information Processing Systems, vol. 29, 2016.

[30] A. Cichocki, R. Zdunek, and S.-i. Amari, “Nonnegative matrix and tensor factorization,” IEEE Signal Processing Magazine, vol. 25, no. 1, pp. 142–145, 2007.

[31] P. O. Hoyer, “Non-negative matrix factorization with sparseness constraints,” Journal of machine learning research, vol. 5, no. Nov, pp. 1457–1469, 2004.

[32] V. P. Pauca, J. Piper, and R. J. Plemmons, “Nonnegative matrix factorization for spectral data aanalysis,” Linear algebra and its applications, vol. 416, no. 1, pp. 29–47, 2006.

[33] S. J. Reddi, S. Kale, and S. Kumar, “On the convergence of adam and beyond,” arXiv Preprint 1904.09237, 2019.

[34] S. Eswar, K. Hayashi, G. Ballard, R. Kannan, M. A. Matheson, and H. Park, “Planc: Parallel low-rank approximation with nonnegativity constraints,” ACM Transactions on Mathematical Software (TOMS), vol. 47, no. 3, pp. 1–37, 2021.

[35] J.-F. Cai, E. J. Candés, and Z. Shen, “A singular value thresholding algorithm for matrix completion,” SIAM Journal on optimization, vol. 20, no. 4, pp. 1956–1982, 2010.

[36] J. W. Demmel, “Superlu users’ guide version 2.0,” University of California at Berkeley, Tech. Rep. UCB/ERL M99/09, 1999. [Online]. Available: http://crd-legacy.lbl.gov/~xiaoye/SuperLU

[37] C. Zheng, M. Yu, J. Shan, A. Wang, and H. Chen, “Fast sparse non-negative least squares via admm for high-resolution doa estimation,” IEEE Sensors Journal, vol. 23, no. 4, pp. 3901– 3910, 2023.

[38] G. H. Golub and C. F. Van Loan, Matrix Computations. JHU Press, 2013.

[39] A. B. Esselman, F. A. Moser, L. E. Tideman, L. G. Migas, K. V. Djambazova, M. E. Colley, E. L. Pingry, N. H. Patterson, M. A. Farrow, H. Yang et al., “In situ molecular profiles of glomerular cells by integrated imaging mass spectrometry and multiplexed immuno?uorescence microscopy,” Kidney International, vol. 107, no. 2, pp. 332–337, 2025.

[40] S. Jain, L. Pei, J. M. Spraggins, M. Angelo, J. P. Carson, N. Gehlenborg, F. Ginty, J.P. Gonçalves, J. S. Hagood, J. W. Hickey et al., “Advances and prospects for the human biomolecular atlas program (hubmap),” Nature Cell Biology, vol. 25, no. 8, pp. 1089–1100, 2023.

[41] B. Efron, “Bootstrap methods: Another look at the jackknife,” in Breakthroughs in Statistics: Methodology and Distribution. Springer, 1992, pp. 569–593.

[42] D. N. Politis and J. P. Romano, “Large sample confidence regions based on subsamples under minimal assumptions,” The Annals of Statistics, pp. 2031–2050, 1994.

[43] G. Casalino, N. Del Buono, and C. Mencar, “Subtractive clustering for seeding non-negative matrix factorizations,” Information Sciences, vol. 257, pp. 369–387, 2014.

