## Appendices 1 for "Unmixing of Imaging Mass Spectrometry Measurements Using Microscopy-Informed Constraints"

### Appendix

#### A. Synthetic Data Generation

The procedure to generate synthetic data is not specific to single cell IMS data. As long as regions of interest are annotated in microscopy or other modality data of high-spatial resolution, this procedure can be used. It comprises three steps:

1) **Spatial View Generation:** A TIFF image containing the high-resolution image with segmented/clustered/classified regions of interest, is processed using the `SPATIAL` module of our `TULIP` toolbox. The TIFF image can be cropped spatially to a certain view and cells that are annotated within this view receive a unique integer IDs. Small cells, *i.e.* only contained in a small number of high-resolution pixels, below a cutoff threshold (*e.g.*, `cut_off`=50 pixels) are removed to reduce the condition number, and the remaining non-cellular regions are designated as non-annotated regions (NARs). Most of the time this cutoff is set to get rid of *e.g.* regions that fall partly inside the view, but only by a few pixels. Finally, the NAR can be subdivided into smaller regions or maintained as a single region.
2) **Spectral View Generation:** Using IMS data, the `SPECTRAL` module extracts spectral signatures by clustering all the spectra. For the cellular component, the spectra are clustered with traditional $k$-means clustering into a fixed number of classes (*e.g.*, `n_clus`=10), while for the background (NAR) a single average spectrum is computed. Therefore, we use only IMS spectra that are fully spanning the background, leaving “mixed” pixels aside. While this might generate a bias towards large cell spectra, we do not consider this aspect as a problem for validation of our method. Another option, is to provide a fixed number of cluster spectra and their probability of appearance, *e.g.* relative cluster sizes.
3) **Synthetic Data Mixing:** The `MIX` module links the spatial and spectral information. Spectral classes are assigned to each cell and NAR via random sampling and a sparse linking matrix. Therefore, we keep the same clustering distribution, *i.e.* keep corresponding cluster sizes. The spatial view is then downsampled (by a factor, *e.g.*, `scale`=16) to mimic the lower resolution of IMS with respect to the microscopy modality used. Optionally, Gaussian-weighted downsampling or other position-specific weights can be applied to emphasize central pixels (at microscopy level) more than pixels (at microscopy level) near the border of the IMS pixel.

Table VI summarizes the key parameters and their roles in the synthetic data generation pipeline. A visual overview of the generation is given in **Fig. 6**.

TABLE VI
Key parameters for synthetic data generation of a synthetic single cell dataset.

| Parameter | Description |
|---|---|
| `cut_off` | Minimum cell size threshold (pixels) |
| `n_clus (cell)` | Number of clusters for cell spectra |
| `n_clus (NAR)` | Number of clusters for NAR spectra |
| `scale` | Downsampling factor (microscopy-to-IMS) |
| `dtype` | Downsampling type (linear/gaussian) |
| `sigma` | Standard deviation for Gaussian kernel |
| `weights` | Positional weights for weighted downsampling |

For our synthetic dataset, we set `cut_off` at 50 pixels, `n_clus (cell)` at 10, `n_clus (NAR)` at 4, `scale` at 16, `dtype` at Gaussian with `sigma` at 4, and no additional `weights`.

#### B. Statistical Resampling Method

To obtain robust estimates of performance metrics and corresponding confidence intervals, we employed a separate bootstrapping procedure for the metrics as well as the spectra.

*1) Metrics:* Specifically, we performed random subsampling (sampling without replacement) of rows and columns from the original data array. In each iteration, a subset of rows was randomly selected, ensuring variability in the input data across repetitions. We used a fixed subset of columns for all iterations. The standard subsample sizes were set to $10,000$ rows and 500 columns per iteration, balancing computational efficiency with statistical robustness. For each subsample, TULIP and the benchmarking methods were applied, and several evaluation metrics were calculated. By repeating this process systematically over multiple iterations, empirical distributions for each metric were obtained. Confidence intervals were subsequently derived using the 0.025 and 0.975 quantiles (CI-0.025 and CI-0.975) to define the lower and upper bounds, respectively, and we report this variation with respect to the median. This method closely aligns with the standard non-parametric bootstrap technique [41], [42], but differs slightly in the use of subsampling rather than full resampling with replacement.

*2) Spectra:* For the generation of confidence intervals on the spectra, bootstrap replicates were generated by resampling the original observations with replacement, thereby preserving the underlying distributional properties of the dataset. Specifically, each bootstrap iteration involved the following two steps:

1) Random selection (with replacement) of indices corresponding to the rows of matrices $A$ and $Y$.
2) Computation of the solution for each bootstrap sample.

This process was repeated for 10 iterations, resulting in a distribution of spectral estimates. The mean and standard deviation across bootstrap replicates provided measures of central tendency and variability, respectively. Confidence intervals were constructed using the normal approximation method, calculated as:

$$\mathrm{CI}_{95\%} = \bar{\theta} \pm 1.96 \times \hat{\sigma}_\theta \tag{13}$$

where $\bar{\theta}$ denotes the mean of bootstrap solutions, and $\hat{\sigma}_\theta$ is their standard deviation.

#### C. HuBMAP Study on Human Kidney (FTU)

The methods (mainly wet-lab specifics) can be found in the original paper [39] and an extended methods section are provided in the supplemental material of the same paper.

#### D. Parameter Tuning

Here, we describe the parameter tuning details per case study. The parameter for NNLS is set heuristically at $\rho = 1$ [37] and number of iterations to 500, and for SVT we set $\delta = 1.2\frac{qp}{\mathrm{nnz}(Y)}$ and $\tau = q$, consistent to advised parameter setting, *e.g.* in [35]. The stopping criteria for SVT are set to be a reconstruction error of $10^{-4}$ or 1000 iterations. To tackle ill-conditioning issues with LS, we use a small regularization term of $\epsilon = 0.001$, that is used in the normal equation as $A^TA + \epsilon I$. For TULIP, we further optimize the parameters of the Adam optimizer and apply an exponential decay on the

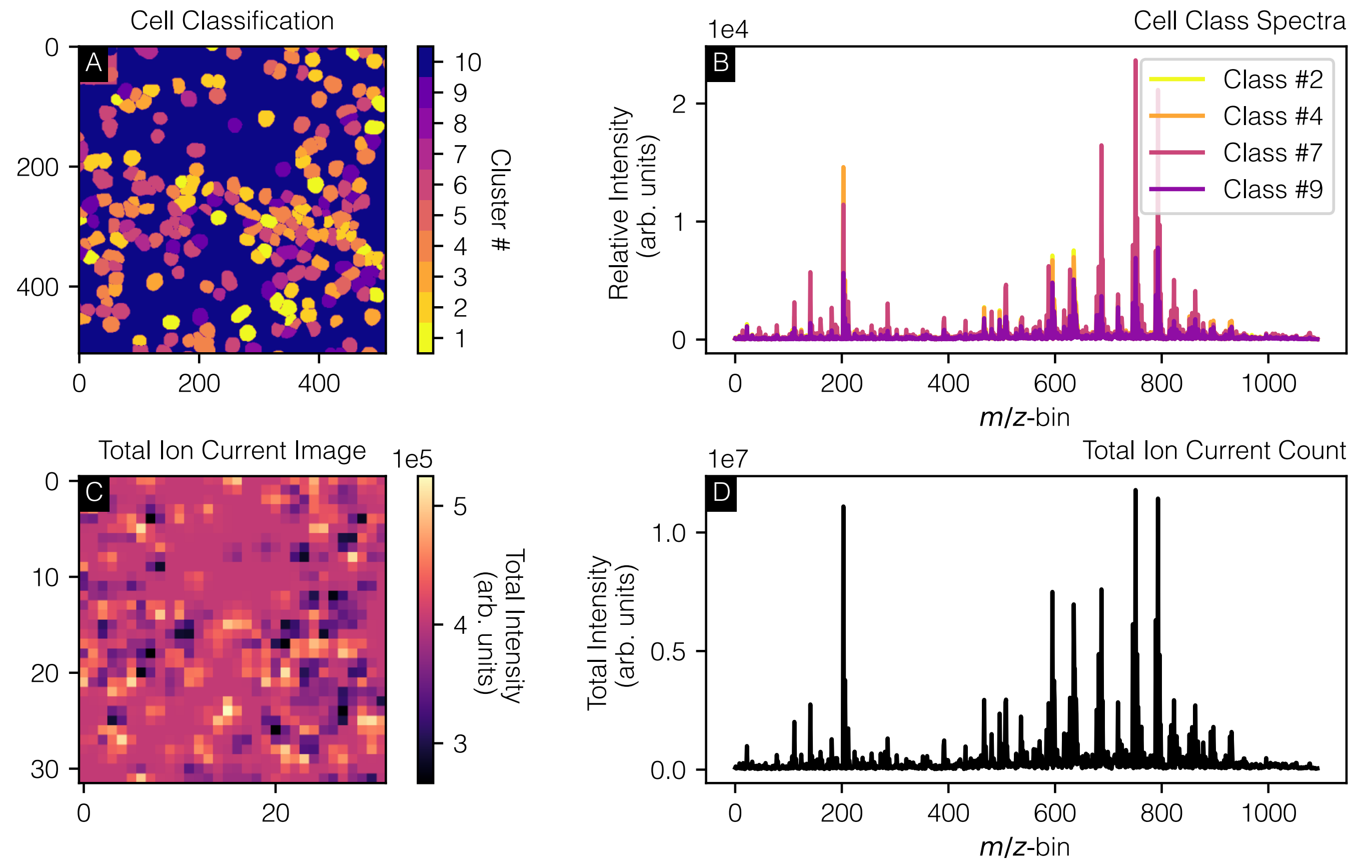


Fig. 6. Synthetic imaging mass spectrometry (IMS) data generation pipeline. (A) A microscopy image is processed to create a spatial map, annotating individual cells and non-annotated regions (NARs), and a cell class is assigned to each cell based on the clustering distribution. (B) Spectral signatures extracted from real IMS data are clustered to define cell classes and a background (NAR) class. These spatial and spectral components are combined through a mixing step, assigning spectral profiles to each spatial region and applying spatial downsampling (linear) to simulate realistic IMS pixel sizes. The resulting synthetic data (C, D) facilitates validation of our unmixing methods for single-cell IMS.

learning rate $\alpha = \alpha_0 \, e^{-r\,k}$, where $\alpha_0$ is specified below, $k$ is the iteration counter and $r$ is a heuristically set decay rate by calculating the halving time to be around $600-700$ steps, leading to $k = 0.001$. Finally, in the first case study we initialize $L$ as a random matrix, with entries sampled from a uniform distribution between 0 and 1 and $R$ through a random *Xcol* initialization [43]. In the second case study, we initialize both $L$ and $R$ as random matrices, with entries sampled from a uniform distribution between 0 and 1 and $R$.

*1) Case Study 1: Overdetermined Scenario:* We set the total number of iterations to 300 and the number of restarts at 20.

*a) Adam Optimizer:* $\beta_1 = 0.9$, $\beta_2 = 0.999$, $\alpha_0 = 2$.

*b) Synthetic Single Cell:* For the synthetic single cell data, the true underlying rank of the problem is known *a priori*, namely 14, hence we fix the rank to that value. For the other parameters, *i.e.* $\lambda_L$, $\lambda_R$ and, $\gamma$ we used a Bayesian optimization with Gaussian process based minimization through `scikit-optimize`. We use a Gaussian kernel and 100 function evaluations. Therefore, we optimize for

$$\theta = \phi_1 + \zeta \left(100 - \bar{\phi}_4(L, R)\right),$$

where $\phi_1$ is the reconstruction score (see Appendix IV-E), $\zeta \in \mathbb{R}$ is a parameter that trades-off sparsity for reconstruction error, and $\bar{\phi}_4(L, R)$ is defined by

$$\bar{\phi}_4(L, R) = \frac{\phi_4(L) + \phi_4(R)}{2},$$

where $\phi_4$ is the sparsity metric (see Appendix IV-E). We set $\lambda_L = [0, 1]$, $\lambda_R = [0, 1]$, $\gamma = [0.1, 1000]$, $\zeta = 0.1$ and optimize $\theta$. We found the following optimal parameters for peak picked data: $\lambda_L^* = 0.1127$, $\lambda_R^* = 0.1380$ and $\lambda^* = 119.0036$. These values were obtained on a separate training set consisting of a different spatial location on the microscopy lay out and a different set of spectra.

*2) Case Study 1: Underdetermined Scenario:* We set the total number of iterations to 300 and the number of restarts to 20.

*a) Adam Optimizer:* $\beta_1 = 0.9$, $\beta_2 = 0.999$, $\alpha_0 = 2$.

*b) Synthetic Single Cell:* For the synthetic single cell data, the true underlying rank of the problem is known *a priori*, namely 14 (10 different cell classes and 4 different background classes), hence we fix the rank to that value. For the other parameters, *i.e.* $\lambda_L$, $\lambda_R$ and, $\gamma$ we used the same Bayesian optimization with Gaussian process as in the previous scenario. We found the following optimal parameters for peak picked data: $\lambda_L^* = 0$, $\lambda_R^* = 0.4211$ and $\lambda^* = 1000.000$. For $\lambda^*$, we noted that increasing the upper bound of its range did not change results, hence, we kept this boundary value. These values were obtained on a separate training set consisting of a different spatial location on the microscopy lay-out and different set of spectra.

*3) HuBMAP Study on Human Kidney (FTU):* For the HuBMAP data, we use a similar method as for the synthetic single cell. However, the rank is here not known *a priori*, and thus becomes an extra parameter to tune. Secondly, given the size of the problem, we train on a single glomerulus (this limits the number of pixels to be below $10,000$) and also subsample the spectral axis by randomly selecting 500 $m/z$-bins. These $m/z$-bins were manually checked to represent high and low intensity peaks. We employ the same Bayesian optimization strategy and obtain the following optimal parameters: $t^* = 100$, $\lambda_L^* = 0$, $\lambda_R^* = 0.3147$ and $\lambda^* = 0.001$. In this case study, we again hit the limits of the search space values for $t$. Heuristically, we found that increasing the rank further only slows the process of finding solutions and does not decrease the reconstruction error substantially. We limited the number of iterations to 100 as on average a step takes around 37 minutes to complete.

*a) Adam Optimizer:* $\beta_1 = 0.9$, $\beta_2 = 0.999$, $\alpha_0 = 2$ and a decay rate $k = 0.01$.

##### E. Metrics

Here, we describe the metrics used for evaluation.

*1) Reconstruction Error:* The reconstruction score ($\phi_1$) measures the accuracy of the unmixing reconstruction, calculated as the relative error between the reconstructed signal and the original mixed signal, expressed as a percentage:

$$\phi_1 = 100\% \times \frac{\|Y - AX\|_F}{\|Y\|_F},$$

where $Y$ contains the IMS spectra, $A$ is the mixing matrix, and $X$ contains the individual estimated spectra of the ROI. In case we are dealing with missing values, we consider the reconstruction error to be defined as:

$$\phi_1 = 100\% \times \frac{\|\mathcal{P}_\Omega(Y - AX)\|_F}{\|\mathcal{P}_\Omega(Y)\|_F}.$$

*2) Cell Fit Score:* Fit quality measures how accurately the cell spectra are reconstructed. Specifically, the cell fit percentage is calculated using the Frobenius norm:

$$\phi_2 = 100\% - 100\% \times \frac{\|X - \hat{X}\|_F}{\|X\|_F},$$

where $X$ contains the ground truth cell spectra and $\hat{X}$ represents the estimated spectra from unmixing.

*3) Non-Negativity:* This metric reports the fraction of non-negative entries in the unmixed spectra matrix $X$, expressed as a percentage:

$$\phi_3 = 100\% \times \frac{\text{number of entries } \geq 0}{\text{total number of entries}}.$$

*4) Sparsity:* Sparsity measures the proportion of nearly-zero entries (less than $2\varepsilon$, where $\varepsilon$ is the machine precision for float32 format) in the estimated unmixed spectra, representing how sparse the recovered solution is:

$$\phi_4 = 100\% \times \frac{\text{number of entries } < 2\varepsilon}{\text{total number of entries}}.$$

*5) Clustering Score:* This metric evaluates the accuracy of clustering after unmixing, calculated as the percentage of correctly matched clusters:

$$\phi_5 = 100\% \times \frac{\text{number of correctly matched clusters}}{\text{total number of clusters}}.$$

A simple $k$-means clustering is applied, from the `sklearn.cluster` module, with the known number of clusters, *i.e.* cell types. We then look at the number of matches of closest clusters making use of the "true" cluster centres and memberships.

*6) Run Time:* The run time is the duration for running of the algorithm. It is given as:

$$\phi_6 = t_{\text{end}} - t_{\text{start}}.$$

##### F. HuBMAP Study on Human Kidney (FTU) - Additional Figures

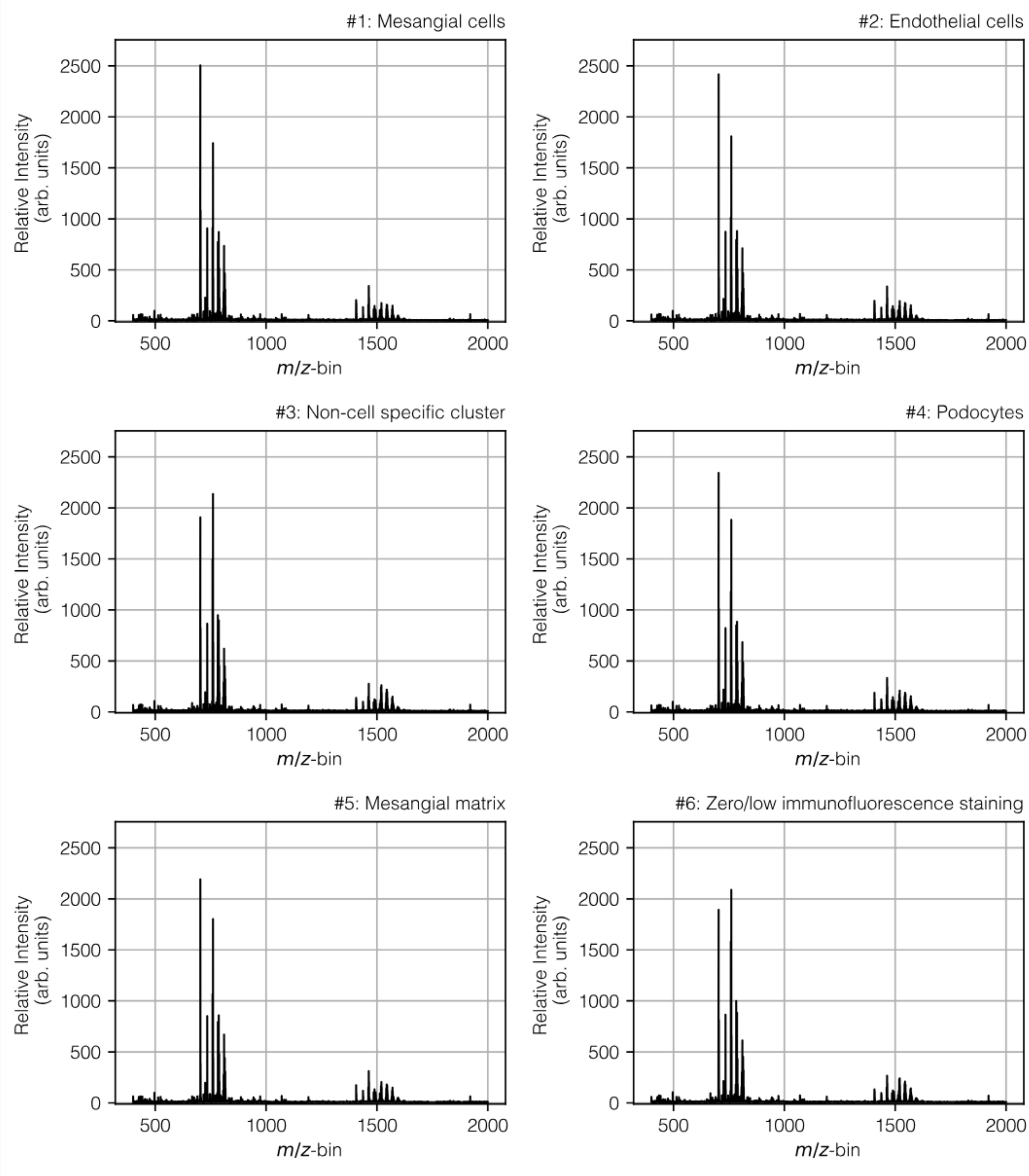


Fig. 7. Estimated spectra for the different glomerular segments. While visually not immediately apparent, all spectra show distinct patterns, *e.g.*, in the region between $500$ and $1000$, we can observe different peaks and intensities.

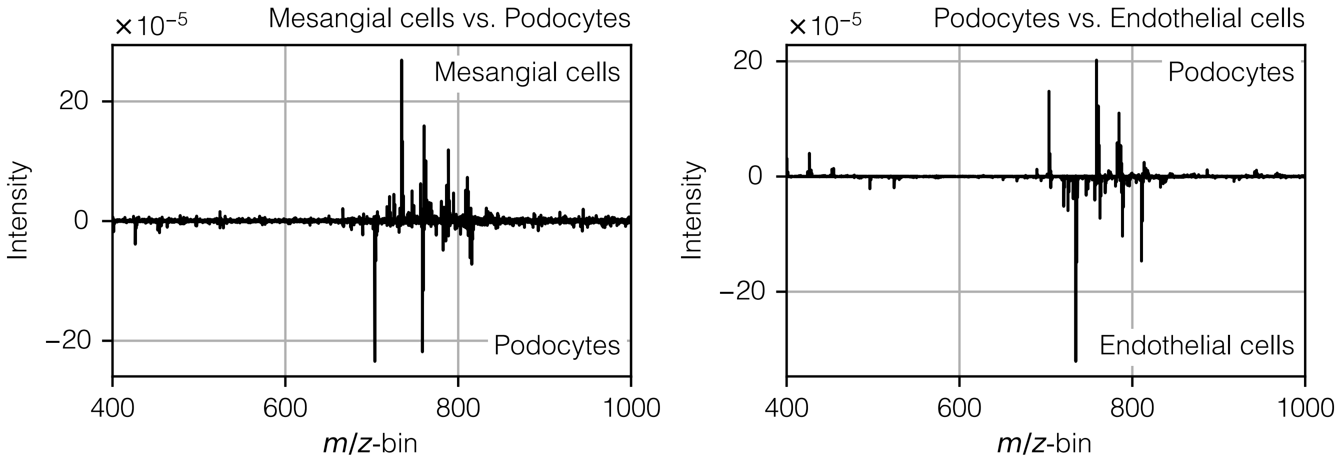


Fig. 8. Mass spectral difference (of total sum normalized spectra) plots illustrating ion intensity differences between ROI $\#1$ (primarily mesangial cells) and ROI $\#4$ (primarily podocytes) (left), and between ROI $\#4$ and ROI $\#2$ (primarily endothelial cells) (right) as obtained in a previous study [39]. These results highlight that each glomerular segment exhibits a distinct profile of IMS-detected molecular species.

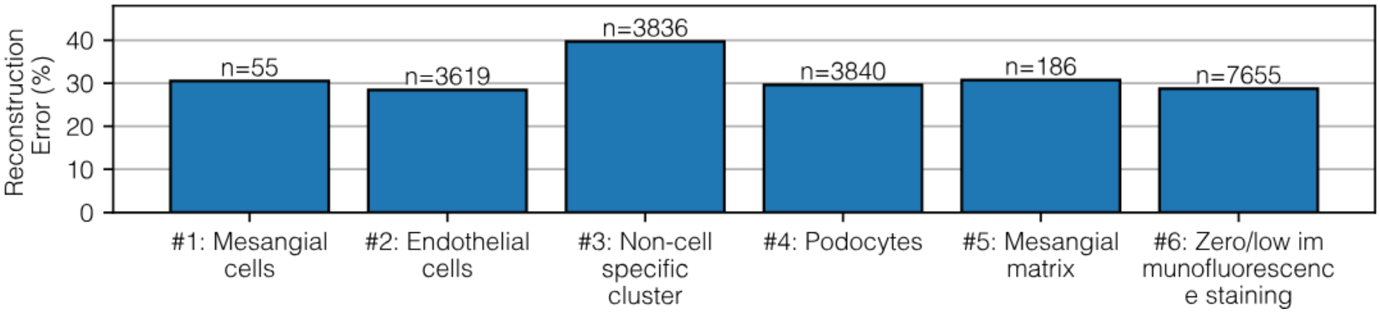


Fig. 9. Bar plot of reconstruction error (%) across six clusters. Each bar represents the percentage error in Frobenius norm, with the sample size $n$ displayed above each bar. The plot highlights how reconstruction accuracy varies by cluster. The reconstruction error is only considered on the pure pixels, only containing a single cluster. Cluster $\mathbf{3}$ (non-cell specific) comprises primarily non-targeted (with the markers) regions within the glomerulus, often capturing a heterogeneous mix of substructures, resulting in a relatively elevated reconstruction error.
